# A novel mechanism of regulation of the transcription factor GLI3 by toll-like receptor signaling

**DOI:** 10.1101/2021.01.01.424866

**Authors:** Stephan J. Matissek, Weiguo Han, Adam Hage, Mona Karbalivand, Ricardo Rajsbaum, Sherine F. Elsawa

**Affiliations:** Deparment of Molecular, Cellular and Biomedical Sciences, University of New Hampshire, Durham, NH; Deparment of Microbiology and Immunology, University of Texas Medical Branch, Galveston, TX; Institute for Human Infections and Immunity, University of Texas Medical Branch, Galveston, TX

**Author notes:** Address correspondence to: Sherine F. Elsawa, PhD 46 College Rd, Rudman 291, Durham, NH 03824.

## Abstract

The transcription factor GLI3 is a member of the GLI family and has been shown to be regulated by canonical hedgehog (HH) signaling through smoothened (SMO). Little is known about SMO-independent regulation of GLI3. Here, we identify TLR signaling as a novel pathway regulating GLI3 expression. We show that GLI3 expression is induced by LPS/TLR4 in human monocyte cell lines and peripheral blood CD14^+^ cells. Further analysis identified TRIF, but not MyD88, signaling as the adapter used by TLR4 to regulate GLI3. Using pharmacological and genetic tools, we identified IRF3 as the transcription factor regulating GLI3 downstream of TRIF. Furthermore, using additional TLR ligands that signal exclusively through TRIF such as the TLR4 ligand, MPLA and the TLR3 ligand, Poly(I:C), we confirm the role of TRIF-IRF3 in the regulation of GLI3. We found that IRF3 directly binds to the GLI3 promoter region and this binding was increased upon stimulation of TRIF-IRF3 with Poly(I:C). Furthermore, using IRF3^−/−^ MEFs, we found that Poly(I:C) stimulation no longer induced GLI3 expression. Finally, using macrophages from mice lacking Gli3 expression in myeloid cells (*M-Gli3^−/−^*), we found that in the absence of Gli3, LPS stimulated macrophages secrete less CCL2 and TNF-α compared with macrophages from wild-type (*WT)* mice. Taken together, these results identify a novel TLR-TRIF-IRF3 pathway that regulates the expression of GLI3 that regulates inflammatory cytokines and expands our understanding of the non-canonical signaling pathways involved in the regulation of GLI transcription factors.

## 1. INTRODUCTION

Hedgehog (HH) signaling is well known for its role in embryonic development, cancer and inflammation (1–4). At the center of HH signaling are the 2 receptors patched (PTCH1) and smoothened (SMO) along with GLI transcription factors (5). In the absence of HH ligand, PTCH1 inhibits SMO. Upon ligand binding to PTCH1, the inhibition of SMO is removed and SMO transduces signal ultimately leading to the activation of GLI proteins (5). Multiple studies have demonstrated that SMO-independent (non-canonical) pathways can regulate GLI proteins (6–10), leading to inflammation and cancer cell growth. Therefore, there is a need to understand the molecular pathways that regulate GLI proteins independent of SMO signaling.

There are 3 members of the GLI family of transcription factors (TFs): GLI1, GLI2 and GLI3. While GLI1 and GLI2 act primarily as transcriptional activators of SMO-dependent HH signaling, GLI3 is known for its transcriptional inhibitory effect where it negatively regulates HH signaling (11–13). Despite the identification of several SMO-independent pathways that regulate GLI1 and GLI2 (10,14), little is known about SMO-independent pathways that regulate GLI3. GLI3 is known to play a role in development-associated diseases such as Greig cephalopolysyndactyly (GCPS) and Pallister-Hall Syndrome (PHS) (15–18). More recent studies suggest a role for GLI3 in the immune system (19).

Immune cells, such as monocytes, recognize pathogens through their pattern recognition receptors (PRR). Toll-like receptors (TLR) are PRR known to be strong inducers of inflammation and immune clearance (20,21). Of the TLRs, TLR4 is known to bind Lipopolysaccharide (LPS), which is expressed on the outer membrane of gram-negative bacteria. Upon LPS binding to TLR4, downstream signaling occurs through either MyD88 or TRIF adapter molecules (22). Signaling through MyD88 activates NF-κB and MAPK, while signaling through TRIF activates IRF3 in addition to MAPK and NF-κB (23–25). All other TLRs signal through MyD88 except TLR3, which signals through TRIF (26). Using either adapter molecule, activation of these transcription factors leads to their nuclear translocation where they regulate the expression of inflammatory genes (27,28)

Here, we identify GLI3 as a downstream target of TLR signaling in monocytes. Stimulation of TLR4 with LPS induces an increase in GLI3 mRNA and protein expression independent of signaling through SMO. Further analysis identified TRIF as the adapter molecule used by TLR4 to modulate GLI3 expression. Using pharmacological inhibitors to target signaling molecules downstream of TRIF, we identify IRF3 as the transcription factor mediating TLR-TRIF-induced GLI3. This was further validated using the TLR4 ligand monophosphoryl lipid A (MPLA) and the TLR3 ligand polyinosinic:polycytidylic acid (Poly(I:C)), which exclusively activate TRIF signaling (29). Both MPLA and Poly(I:C) increased GLI3 expression and overexpression of IRF3 increased GLI3 expression. We found that IRF3 directly binds the GLI3 promoter region upstream of exon 1 and this binding was increased upon activation of TRIF-IRF3 signaling. Using mouse embryonic fibroblasts (MEFs) lacking IRF3 expression (IRF3^−/−^ MEFs), we show that activation of TRIF-IRF3 signaling was no longer able to increase GLI3. Finally, using macrophages from mice lacking Gli3 expression in myeloid cells (*M-Gli3^−/−^*), we found that LPS stimulation results in significantly reduced CCL2 and TNF-α levels compared with macrophages from WT mice. Taken together, this study identifies TLR-TRIF-IRF3 signaling as a novel pathway that regulates GLI3 expression and highlights a novel role for Gli3 in regulating inflammatory cytokines in response to LPS-TLR4 signaling.

## 2. MATERIALS AND METHODS

### 2.1 Cell lines and primary cells

The cell line Mono-Mac-6 (MM6) was purchased from DSMZ and cultured in RPMI 1640 supplemented with 10% FBS, 1% 2mM L-glutamine, non-essential amino acids, 1mM Sodium Pyruvate, and 10μg/ml human insulin. THP-1 and U937 cell lines were purchased from ATCC. THP-1 cells were cultured in RPMI 1640 with 10 % FBS supplemented with 0.05 mM β-Mercaptoethanol and U937 cells were grown in RPMI 1640 with 10% FBS. An antibiotic-antimycotic was added to all cell culture media. Wild-type (WT) and *IRF3*^−/−^ MEFs were a generous gift from Benjamin R. tenOever, and were generated as previously described (30). MEFs were maintained in Dulbecco’s Modified Eagle’s Medium (DMEM) (Corning, NY) supplemented with 10% v/v fetal bovine serum (FBS) (HyClone) and 1% v/v penicillin-streptomycin (Corning). Poly(I:C) HMW stimulations (10 μg/mL) were performed with Lipofectamine 2000 according to the manufacturer recommendations using a ratio of 1 μg:0.2 μL (Poly(I:C):Lipofectamine 2000).

Primary human monocytes were purified from peripheral blood mononuclear cells (PBMCs) obtained from buffy coats purchased from the blood bank of New York (NYC, NY) or the Oklahoma Blood Institute (Oklahoma City, OK). Experiments using cells from different donors are numbered D1-D6. Monocytes were isolated by negative selection using the monocyte enrichment kit from STEMCELL Technologies (Cambridge, MA). Cell purity was confirmed by flow cytometry using CD14+ staining and was found to be >90%. Primary cells (CD14+) were plated and treated immediately after purification as described.

### 2.2 Reagents

Lipopolysaccharide (LPS) (*E.coli, Serotype 0111;B4*) and Poly(I:C) were purchased from MilliporeSigma (St. Louis, MO). CpG was purchased from Integrated DNA Technologies (Coralville, IO). The TLR4 signaling inhibitor CLI-095) and MPLA were purchased from Invivogen (San Diego, CA). The MAPK inhibitor (PD98059), NF-κB inhibitor (QNZ), TBK1 inhibitor (BX-795) and SMO inhibitor (Cyclopamine) were purchased from SelleckChem (Houston, TX).

### 2.3 RNA isolation and Quantitative RT-PCR (qPCR)

For total RNA isolation, TRIsure reagent (Bioline, obtained through Thomas Scientific, Swedensboro, NJ) was used following the manufacturer’s recommendations as previously published (31,32). cDNA synthesis was performed using Moloney Murine Leukemia Virus Reverse Transcriptase (MML-V)(Promega Corporation, Madison, WI). ViiA-7 instrument (Life Technologies, Grand Island, NY) was used to perform qPCR analysis. Amplification results from GLI1, GLI2 and GLI3 were normalized to human GAPDH. The following qPCR primers were used GAPDH, 5’-CTCGACTTCAACAGCGACA-3’ (forward) and 5’-GTAGCCAAATTCGTTGTCATACC-3’ (reverse). GLI1, 5’-TCACGCCTCGAAAACCTGAA-3’ (forward) and 5’-AGCTTACATACATACGGCTTCTCAT-3’ (reverse); GLI2, 5’-GGAGAAGCCATATGTGTGTGAG-3’ (forward) and 5’-CAGATGTAGGGTTTCTCGTTGG-3’ (reverse); GLI3, 5’-GGGACCAAATGGATGGAGCA-3’ (forward) and 5’-TGGACTGTGTGCCATTTCCT-3’ (reverse).

### 2.4 Chromatin immunoprecipitation (ChIP) assay

Briefly, 10×10^6^ cells were cross-linked using 0.37% formaldehyde for 10 min at room temperature then lysed and sonicated to shear DNA to ~ 500 bp (QSonica, Q800R3). Samples were immunoprecipitated using anti-IRF3 antibody or isotype control (Biolegend, San Diego, CA) using magnetic Protein A/G beads (ThermoFisher Scientific) at 4°C overnight. Samples were washed and reverse crosslinked overnight in 5M NaCL at 65°C. After RNase A and Proteinase K digestion, DNA was purified using GeneJET PCR Purification Kit (ThermoFisher Scientific) and used for qPCR analysis using the following primers: Binding site 1 (BS1), 5’-GGCTAAGAATGGAGTGTTTGGA-3’ (forward) and 5’-CACCACTGGTTTAGCTACATACAT-3’ (reverse); Binding site 2 & 3 (BS2/3), 5’-CTGGGCAACAGAGTGAGAC-3’ (forward) and 5’-CACCCGGAACCTCAATGTTA-3’ (reverse); Binding site 4 (BS4), 5’-CTGCCCTTGGGACTCAC-3’ (forward) and 5’-CAAAGGGCACTGACAAAGTATT-3’ (reverse). To determine the effect of TLR-TRIF signaling on IRF3-mediated regulation of GLI3, 10×10^6^ cells were cultured at a density of 5×10^6^ cells/ml and treated with 10μg/ml Poly(I:C) for 1h or left alone followed by fixation and ChIP analysis as described earlier.

### 2.5 Plasmid constructs and cell transfections

Short hairpin RNA targeting TRIF and MYD88 (shTRIF, shMYD88) were purchased from OriGene Technologies (Rockville, MD). The plasmid expressing dominant negative TRIF (dnTRIF) was purchased from Invivogen (San Diego, CA). Transfections were performed by electroporation using BTX ECM 630 (Holliston, MA) using the following parameters: MM6: 240v/25ms and U937: 300v/10ms. Five million cells were electroporated with 10μg shTRIF, shMYD88 or scrambled control (shScr) or 5μg dnTRIF or empty vector in OPTI-MEM followed by culture for 48h.

### 2.6 Immunoblotting

Immunoblotting was performed as previously described (31,32). Cell lines and primary monocytes (5×10^6^ cells/ml) were treated as indicated and harvested in 100 μl RIPA buffer (ThermoFisher Scientific). Total protein concentration was determined using a BCA assay (ThermoFisher Scientific). For GLI3, 5% self-prepared SDS gels were used while for all other proteins 10% gels were used. Transfer to nitrocellulose membranes was performed using a turbo transblot system (Biorad, Hercules, CA). GLI3 antibody was purchased from abcam (cat# ab123495), β-actin antibody was purchased from Milipore Sigma (cat# A3854), IRF3, phospho-IRF3, ERK, phospho-ERK, IκBα, TRIF and MYD88 antibody were purchased from cell signaling (cat#4302, cat# 4947, cat# 9102, cat# 9101, cat# 9242, cat#4596, cat#4283 respectively).

### 2.7 Mice

*Gli3^fl/fl^* mice and lysozyme M (*LysM*)*-cre* mice were obtained from Jackson Labs (Bar Harbor, ME). Mice were bred and maintained at the UNH animal resource office (ARO). Mice were crossed to generate mice with conditional deletion of *Gli3* in myeloid cells (*M-Gli3^−/−^*). Mice were genotyped at p6-p9 using toe clipping and were separated into *M-GLI3^−/−^* or *WT* groups. All experiments were approved by the UNH IACUC following guidelines of the National Institutes of Health. Mice were used for experiments when they reached 8-14 weeks of age.

Genotyping was performed using DNA extracted from mice toes using Phire Animal Tissue Dircet PCR Kit (Fisher Scientific, F-140WH). The following PCR primer pairs were used: Gli3 flox, 5’-GATGAATGTGATCCAGGGC-3’ (forward) and 5’-GTCATATTGTGCCCAGTAGTAGC-3’ (reverse); Cre, 5’-GCGGTCTGGCAGTAAAAACTATC-3’ (forward) and 5’-GTGAAACAGCATTGCTGTCACTT-3’ (reverse); and recombined Gli3 (rGLI3), 5’-CTGGATGAACCAAGCTTTCCATC-3’ (forward) and 5’-CAGTAGTAGCCTGGTTACAG-3’ (reverse). Samples were analyzed on 2% agarose gels.

### 2.7 Enzyme-linked immunosorbent assay (ELISA)

Mouse IL-6 ELISA (R & D Systems; DY406), mouse CCL2 ELISA (R & D Systems; DY479) and mouse TNFα ELISA (R & D Systems; DY410) were used to quantify cytokine secretion in cultured peritoneal macrophages. Peritoneal macrophages were isolated by injection of mice with 500 μl incomplete Freund’s Adjuvant (IFA) i.p. then after 3 days, mice were euthanized and macrophages were harvested via peritoneal lavage with IMDM containing 10% FBS as previously described (33). Peritoneal cells were washed once and 1×10^6^/ml were allowed to adhere to cell culture plates for 1h. Cells were then gently washed with DPBS, and 1 ml of fresh media (IMDM+10%FBS+1%A/A) was added with either 100 ng/ml LPS (E.coli; Serotype 0111:B4; Sigma Aldrich) or control for 24h. Supernatants were harvested and diluted 1:40 for then used in ELISA following manufacturer recommendations.

### 2.8 Statistical analysis

Statistical analysis was performed using GraphPad Prism software (San Diego, CA). A two-tailed t test was used to determine statistical significance between two variables and a two-way ANOVA was used to compare more than 2 variables. Statistical significance is indicated on each figure as follows: * (p<0.05), ** (p<0.01), *** (p<0.001), and **** (p<0.0001).

## 3. RESULTS

### 3.1 LPS stimulation induces GLI3 expression

We stimulated human monocyte cell lines MM6, THP-1 and U937 with LPS and investigated the effects on GLI expression. We found that LPS stimulation induces an increase in *GLI3* mRNA expression in all 3 cells lines (Fig. 1A). *GLI1* expression was not altered in cell lines, while *GLI2* expression was only induced in THP-1 cells. Using CD14^+^ cells purified from peripheral blood mononuclear cells (PBMCs), we confirm that LPS stimulation induces *GLI3* expression in primary human monocytes (Fig. 1B). The increase in *GLI3* expression upon LPS stimulation occurred in a time and dose-dependent manner (Fig. 1C). This increase in *GLI3* expression also resulted in an increase in GLI3 protein in response to stimulation with LPS (Fig. 1D). Taken together, these results propose the TLR4 signaling as a novel non-canonical HH pathway that can regulate GLI3 expression.

**Figure 1:**
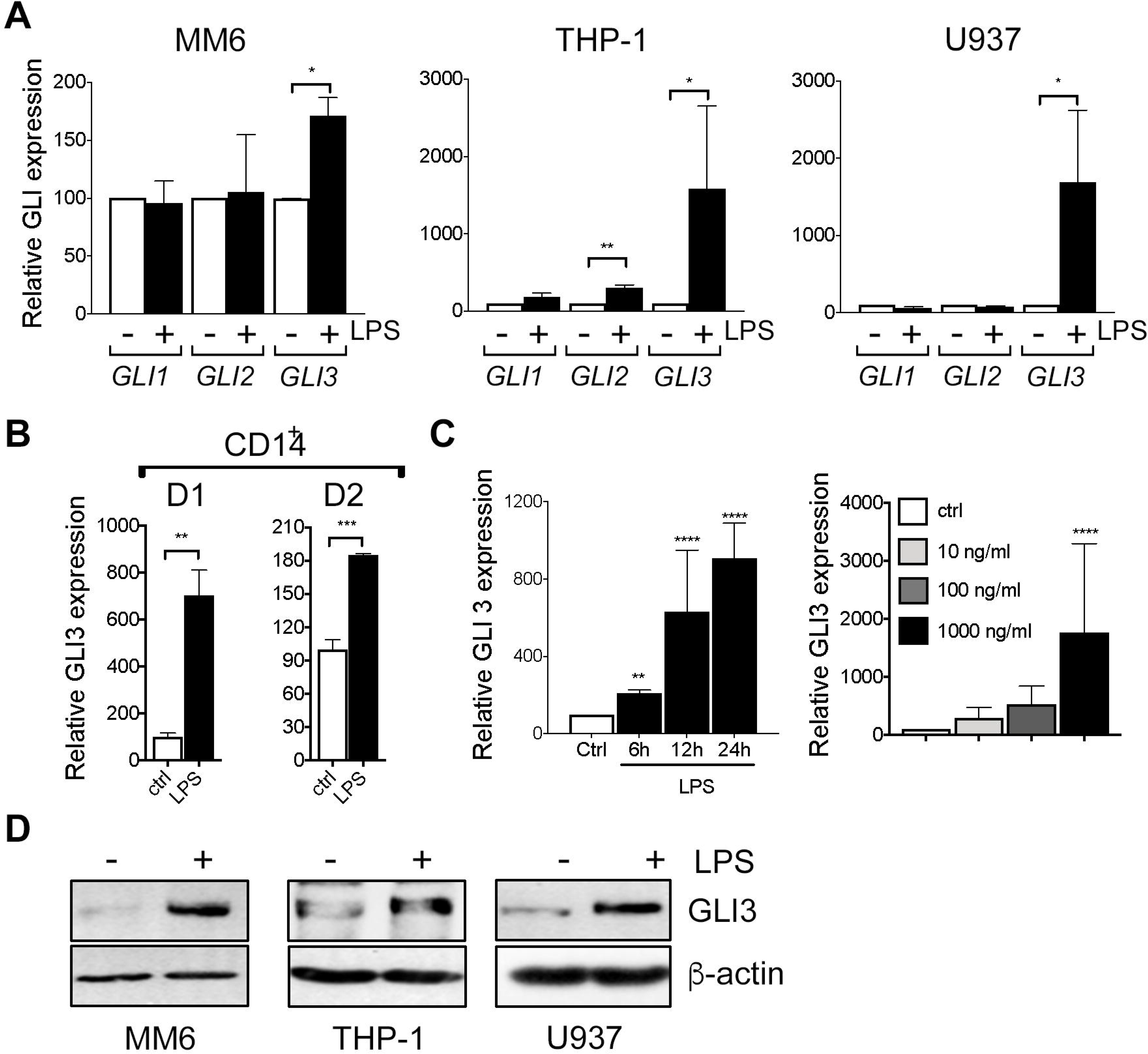
LPS induces GLI3 expression in a HH independent manner. (A) Human monocyte cell lines (MM6, THP-1, U937) (2 × 10^6^ cells/ml) were treated with/without 100 ng/ml LPS (0111; B4 *E.coli*) for 1 h followed by determination of GLI expression by qPCR. (B) CD14^+^ cells from PBMCs (2 × 10^6^ cells/ml) were stimulated with 100 ng/ml LPS for 1 h followed by qPCR to determine GLI3 expression. (C) MM6 cells (2 × 10^6^ cells/ml) were stimulated with 100 ng/ml LPS for the indicated times, or with the indicated doses of LPS for 1 h followed by determination of GLI3 expression by qPCR. (D) Human monocytes (5 × 10^6^ cells/ml) were stimulated with LPS for 12 h followed by determination of GLI3 protein expression by immunoblotting. All experiments were repeated at least three times. Data are presented as average of at least three independent experiments ± SEM.

### 3.2 LPS/TLR4-induced GLI3 occurs through TRIF signaling

To elucidate the signaling mechanism downstream of TLR4 that results in increased GLI3 expression, we used the TLR4 signaling inhibitor (CLI-095), which inhibits TLR4 signaling intracellularly and found that monocyte cell lines treated with the TLR4 signaling inhibitor had reduced GLI3 expression (Fig. 2A). Furthermore, in the presence of the TLR4 signaling inhibitor, LPS was not able to induce GLI3 expression (Fig 2B) suggesting that an active TLR4 signaling pathway was required for LPS-induced GLI3.

**Figure 2:**
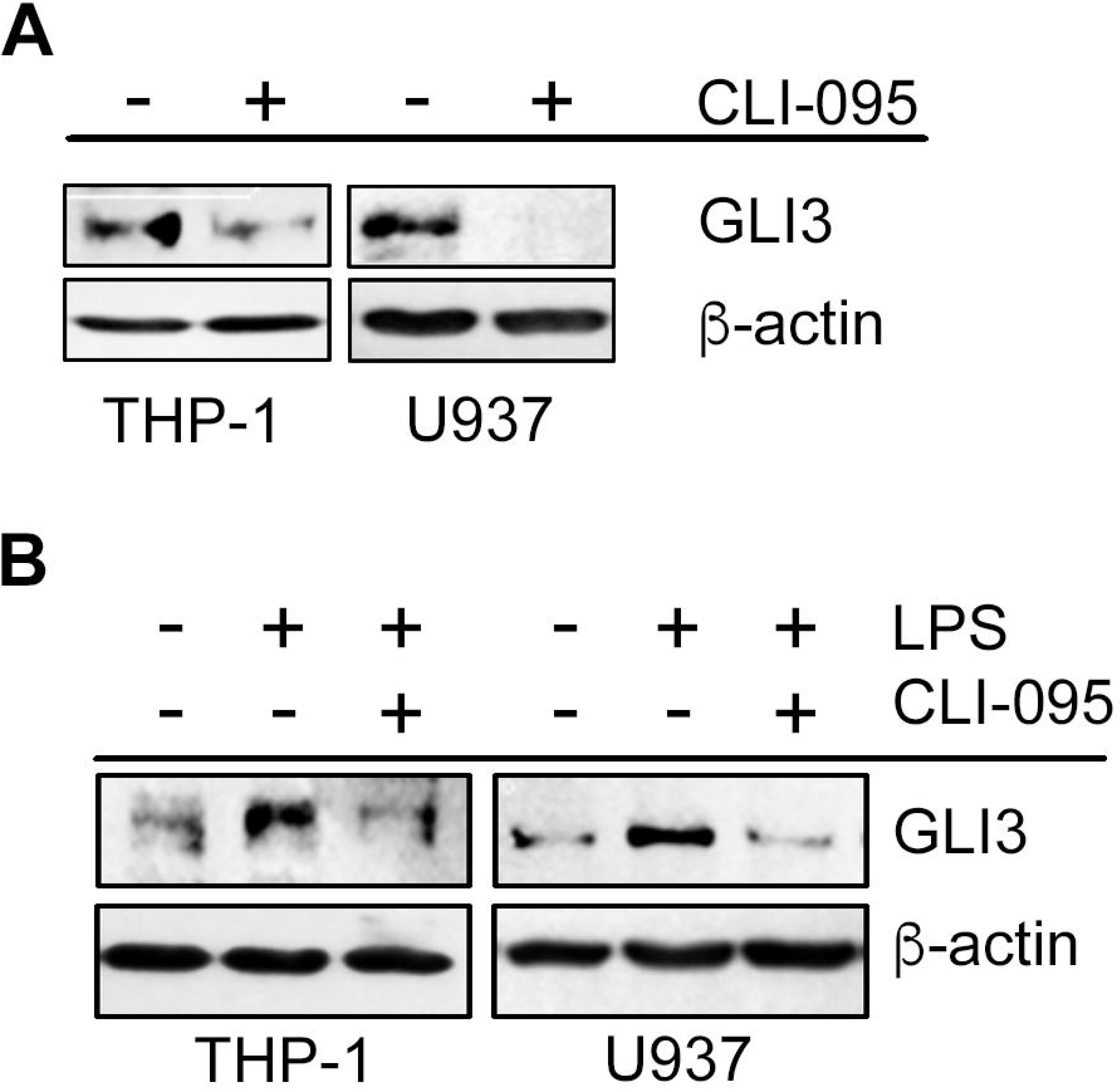
Functional TLR4 is required for GLI3 expression. (A) Monocytes (5×10^6^ cells) were treated with 1 μg/ml TLR4 signaling inhibitor (CLI095) for 12h followed by determination of GLI3 expression by western blot. (B) Monocytes (5×10^6^ cells) were pretreated with CLI095 for 12 h followed by stimulation with 100 ng/ml LPS for an additional 12 h. GLI3 expression was determined by western blot. These experiments were repeated 3 times.

LPS-TLR4 signaling can utilize one of 2 adapter proteins, MyD88 or TRIF (28). To determine the signaling mechanism downstream of TLR4 that regulates GLI3, we used RNAi targeting either MyD88 or TRIF to determine the effect on GLI3 expression. When cells were transfected with RNAi targeting TRIF, but not MyD88, there was a reduction in GLI3 expression (Fig. 3A). To validate this, we utilized a dominant negative form of TRIF (dnTRIF) that lacks N- and C-terminus and harbors a proline to histidine mutation at amino acid 434 which is crucial for TLR-dependent signaling. We found that in cells expressing dnTRIF, there was a reduction in GLI3 protein expression (Fig. 3B). Additionally, LPS stimulation was no longer able to induce GLI3 protein expression in cells transfected with shTRIF (Fig. 3 C). Taken together, these results suggest that LPS-TLR4-induced GLI3 occurs via signaling through TRIF.

**Figure 3:**
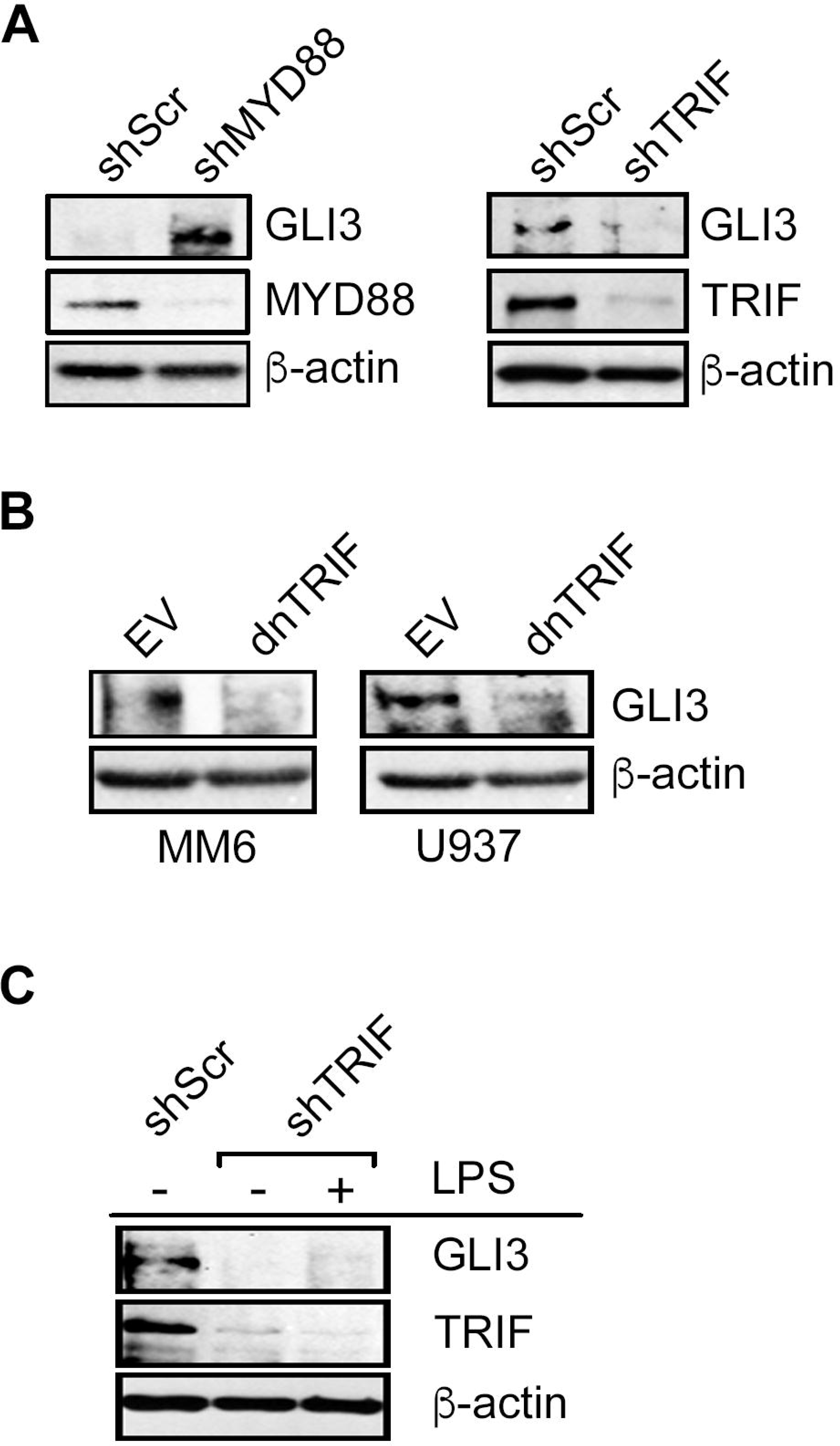
LPS-induced GLI3 is regulated by TRIF downstream of TLR4. (A) U937 cells (10×10^6^ cells) were transfected with 10 μg of either shMYD88, shTRIF or scrambled controls (shScr). After 2 days, cells were lysed and lysates were used to determine protein expression by western blot. (B) Monocytes (10×10^6^ cells) were transfected with a dominant negative form of TRIF (dnTRIF) or empty vector (Ctrl) for 2 days followed by determination of protein expression by western blot. (C) MM6 cells (10×10^6^ cells) were transfected with shTRIF or shScr and cultured for 30 h, followed by treatment with 100ng/ml LPS. After an additional 12 h, cells were lysed and lysates were used for western blot to determine protein expression.

### 3.3 TLR4-TRIF regulates GLI3 through the transcription factor IRF3

Following activation of TRIF, three different downstream signaling pathways can be initiated: NF-κB pathway, MAPK pathway and IRF3 pathway (28). We used pharmacological inhibitors to target each of these pathways to identify the contribution of each in TLR4-TRIF-mediated regulation of GLI3. Western blot analysis shows that only inhibition of IRF3 signaling using the TBK1 inhibitor BX795 reduced basal levels of GLI3 (Fig. 4A). Pretreatment of monocytes with BX795 also resulted in a loss of LPS-induced GLI3 (Fig. 4B). To further validate the role of IRF3 as a regulator of GLI3, we overexpressed IRF3 in U-937 cells and found an increase in GLI3 expression (Fig. 4C). Furthermore, we stimulated monocytes with monophosphoryl lipid A (MPLA), a TLR4 ligand that signals exclusively through TRIF and found an induction in GLI3 expression (Fig. 4D). Finally, we compared GLI3 expression in response to LPS (TLR4 agonist which signals though MyD88 and TRIF), CpG (TLR9 agonist which signals through MyD88) and Poly(I:C) (TLR3 agonist which signals through TRIF) and found a robust increase in GLI3 expression in response to Poly(I:C) in monocyte cell lines and CD14+ cells from PBMCs, further supporting a role for TRIF-IRF3 signaling in the regulation of GLI3 (Fig. 4E). Taken together, these data suggest that GLI3 is a novel target of TRIF-IRF3 signaling and can be regulated by TLR3 and TLR4.

**Figure 4:**
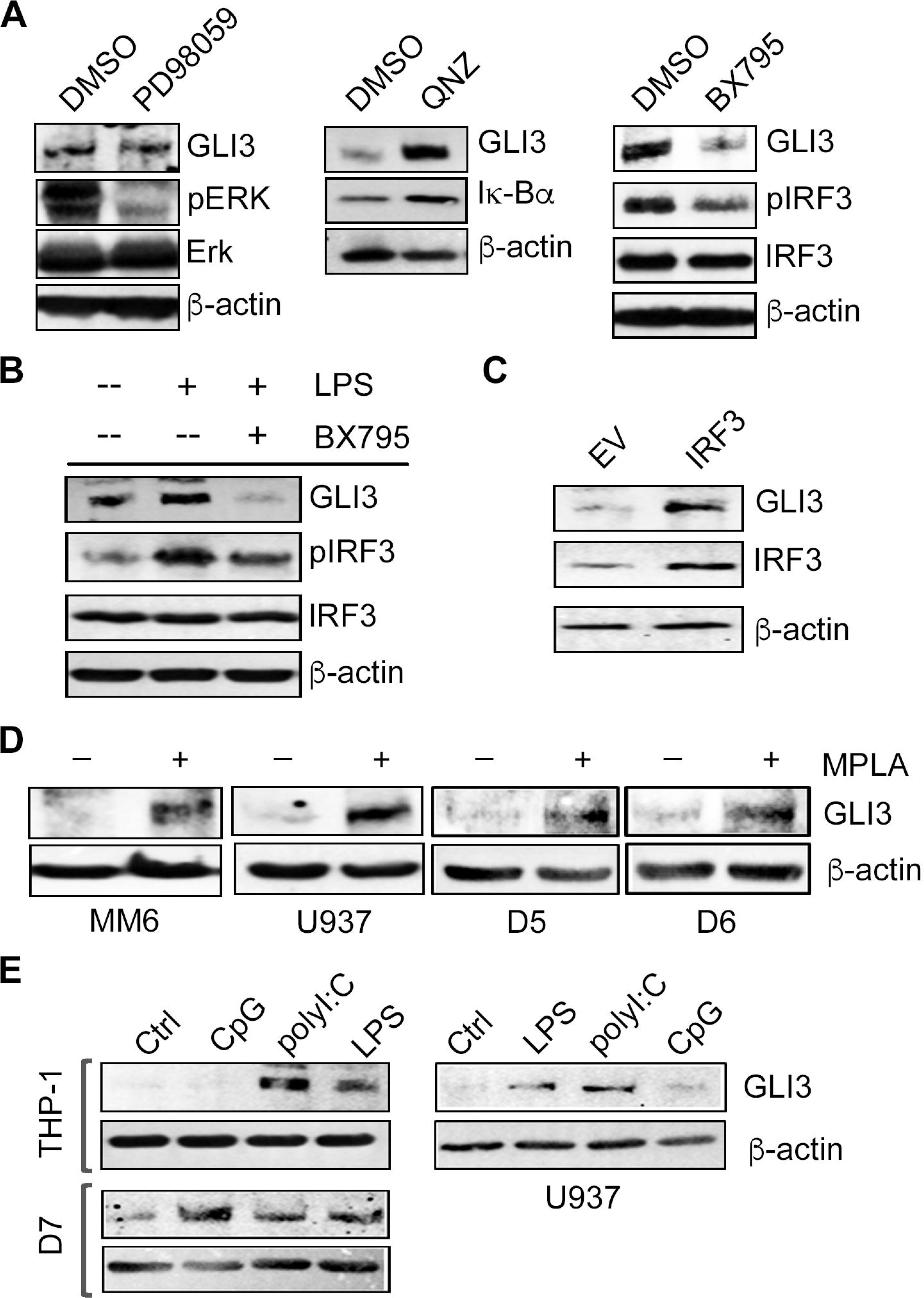
GLI3 is regulated by IRF3 downstream of TRIF. (A) THP-1 cells (5×10^6^ cells) were treated with 50μM Erk inhibitor (PD98059) or DMSO control for 30 min, 500 nM NF-κB inhibitor (QNZ) or DMSO for 2 h, or 1 μM TBK1 inhibitor (BX795) or DMSO for 1 h. Cells were harvested, lysed and lysates were used to determine protein expression by western blot. (B) THP-1 cells (5×10^6^ cells) were pretreated with 1 μM BX795 for 1 h, followed by stimulation with 100 ng/ml LPS for 12h. Cells were lysed and lysates were used to determine protein expression by western blot. (C) U937 cells (10×10^6^ cells) were transfected with an IRF3 expression construct or empty vector (EV) for 2 days followed by lysis and western blot. (D) Monocytes (5×10^6^ cells) were treated with 1 μg/ml TLR4 agonist MPLA for 12 h followed by western blot to determine GLI3 expression. (E) Monocyte cell lines and primary donor CD14^+^ cells (D7) (5×10^6^ cells) were treated with 10 μg/ml CpG, 10 μg/ml Poly(I:C), 100 ng/ml LPS or left untreated (Ctrl) for 12h followed by western blot to determine GLI3 expression. Data shown are representative of three independent experiments.

### 3.4 IRF3 directly binds the GLI3 promoter

Although several studies have characterized the promoter region of GLI1 and GLI2 (34,35), less is known about the GLI3 promoter or it’s regulation. The *GLI3* gene is composed of 15 exons with transcription of the gene beginning in exon 2 (19). Therefore, we examined approximately 3000 bp upstream of ATG of the GLI3 gene for candidate IRF3 binding sequences and identified 4 consensus sequences (AANNGAAA) that IRF3 may bind to regulate GLI3 expression (Fig. 5A). Using MM6 cells (which have the highest copy number of GLI3 [data not shown]), we performed chromatin immunoprecipitation (ChIP) assay to determine IRF3 binding to the GLI3 promoter region and found IRF3 binding to all candidate sites (Fig. 5B). Activation of TRIF signaling in MM6 cells using Poly(I:C) resulted in a significant increase in IRF3 binding to all candidate IRF3 binding sites (Fig. 5C). Furthermore, using CD14^+^ cells purified from PBMCs that were treated with Poly(I:C), there was a significant increase in IRF3 binding to the GLI3 promoter in response to Poly(I:C) stimulation (Fig. 5D); and stimulation of CD14^+^ cells from the same donor with Poly(I:C) increases GLI3 expression (Fig. 5E). Taken together, these results suggest that TLR-TRIF-mediated regulation of GLI3 occurs via direct binding of IRF3 to the GLI3 promoter and identifies IRF3 as a novel transcription factor that regulates GLI3 independent of HH signaling.

**Figure 5:**
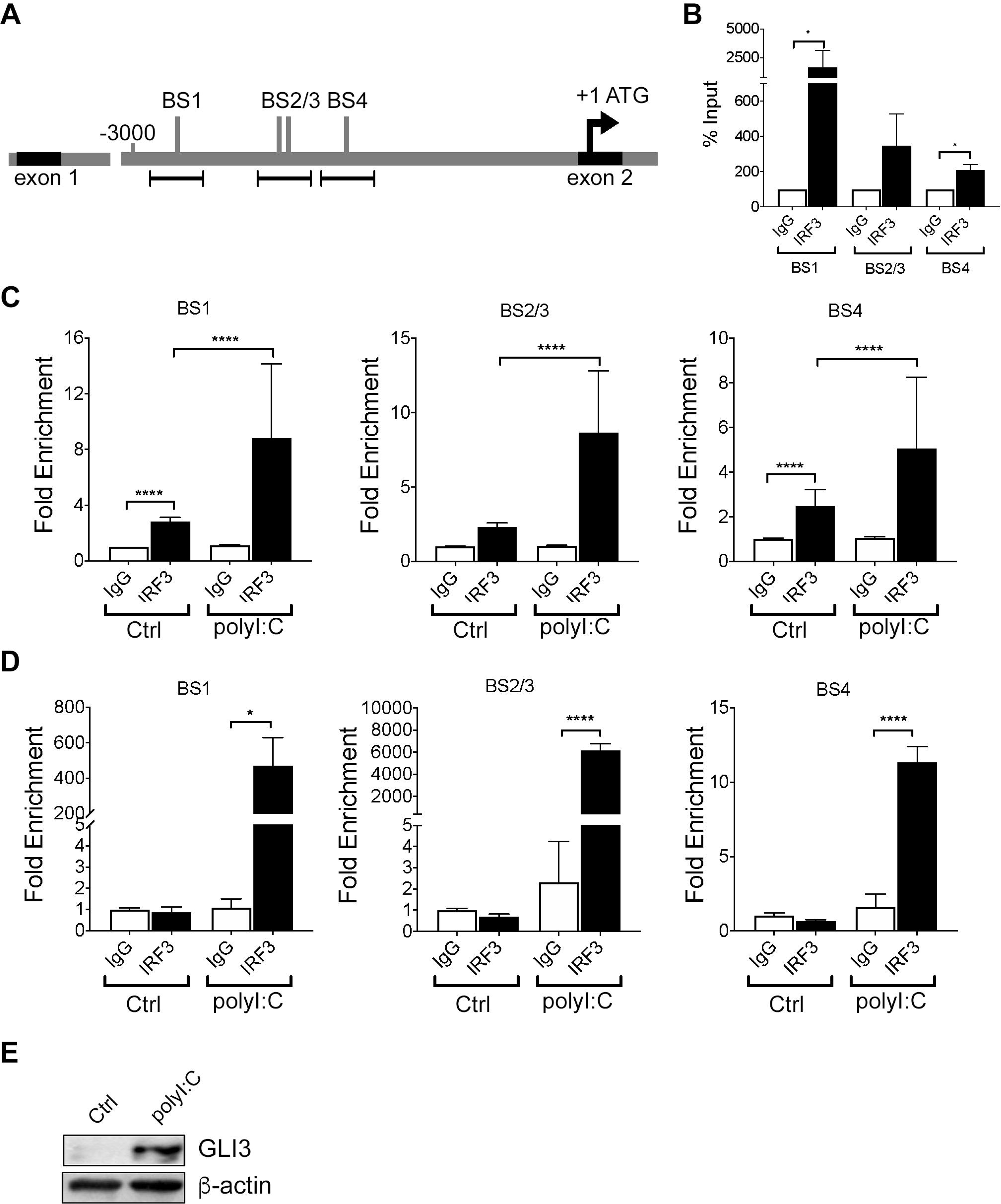
IRF3 regulates GLI3 by directly binding to IRF3 binding sites. (A) Schematic diagram of ~ 3000 bp upstream of ATG located in exon 2 of GLI3. Several candidate IRF3 binding sites (BS) were identified. (B) Untreated MM6 cells (10×10^6^) were lysed and a chromatin immunoprecipitation (ChIP) assay was performed followed by qPCR using three primer sets for the areas containing IRF3 binding sites upstream of ATG of GLI3.(C) MM6 cells (10×10^6^) were treated with 10μg/ml Poly(I:C) for 1h prior to ChIP and qPCR.(D) Primary human monocytes (D3) were treated with 10μg/ml Poly(I:C) for 1h prior to ChIP and qPCR. (E) Primary human monocytes (D3) were challenged with 10μg/ml Poly(I:C) for 12h prior to western blot analysis.

### 3.5 IRF3 is required for TRIF-induced GLI3

To confirm the role of IRF3 in TLR-TRIF-induced GLI3, we used mouse embryonic fibroblasts (MEFs) that lack IRF3 (*IRF3^−/−^*) or wild-type (WT) MEFs stimulated with Poly(I:C). Consistent with previous results in monocytes, in WT MEFs treated with Poly(I:C), there was a time-dependent increase in Gli3 expression (Fig. 6A). This Poly(I:C)-induced Gli3 was lost in *IRF3^−/−^* MEFs. This finding was consistent with an increase in interferon beta (IFN-β) expression in the WT MEFs, but not in the *IRF3^−/−^* MEFs (Fig. 6A). Taken together, these results support the requirement for IRF3 in TLR-TRIF-induced *Gli3*.

**Figure 6:**
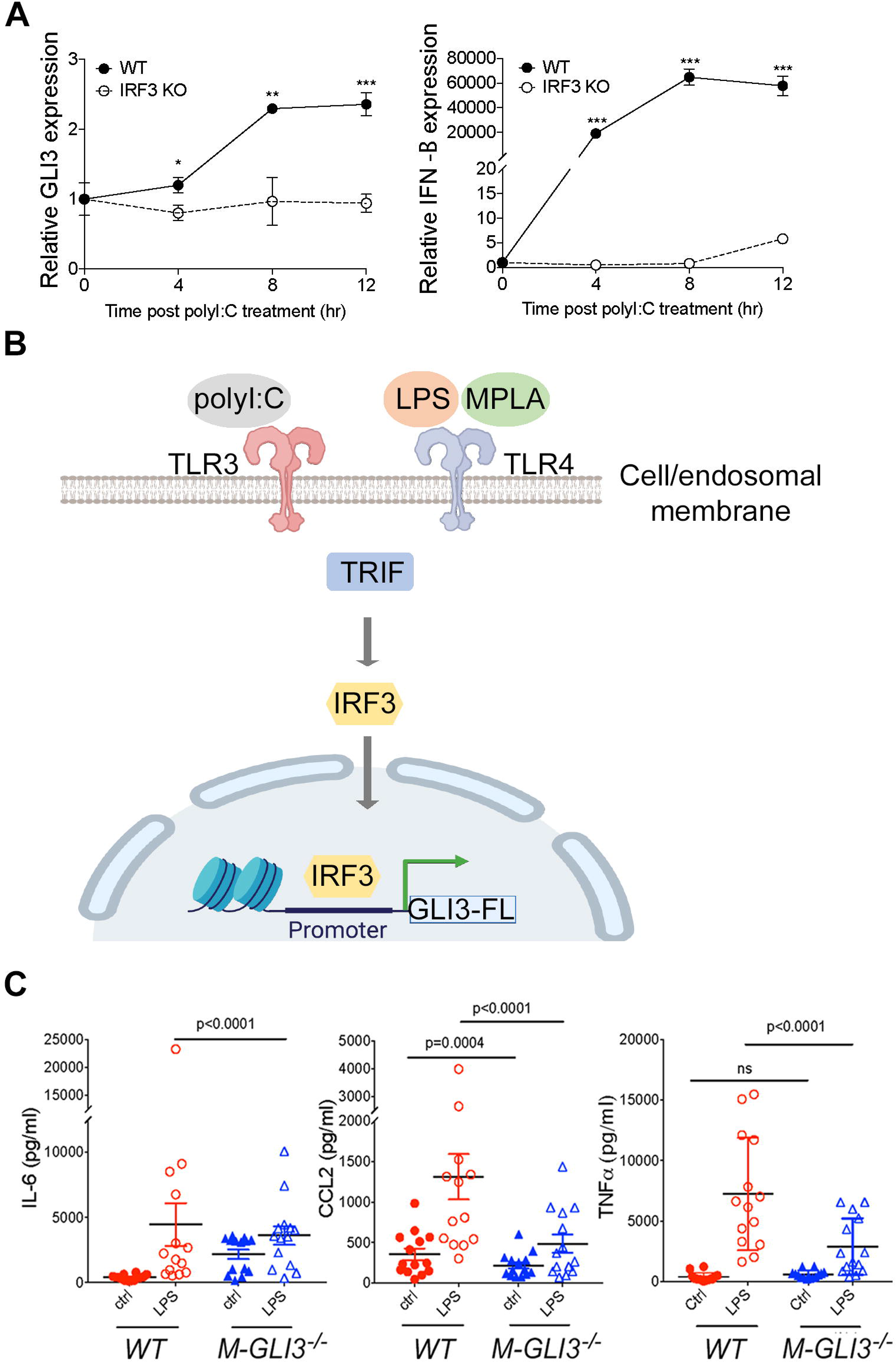
Gli3 mediates TLR-induced inflammation. (A) *WT* and *IRF3^−/−^* MEFs (1.5×10^5^) were treated with 10 μg/mL Poly(I:C) for the indicated time followed by qPCR to determine GLI3 expression. (B) Model of TLR-TRIF-IRF3-dependent regulation of GLI3. (C) IFA-elicited macrophages from *M-Gli3^−/−^* or WT mice (n=14/group) were cultured in the presence or absence of 100 ng/ml LPS for 24 hours. Supernatants were used to quantify cytokine levels by ELISA.

### 3.6 Gli3 mediates TLR-TRIF-induced inflammation

Because GLI3 was induced in response to TLR stimulation, we hypothesized that GLI3 regulates inflammatory genes expressed in response to TLR signaling. Since Gli3^−/−^ mice are embryonically lethal(36,37), we generated mice with conditional Gli3 knockout in myeloid cells (*M-Gli3^−/−^*) by crossing *Gli3^fl/fl^* mice with *LysM-cre* mice. We verified the lack of Gli3 expression by PCR by confirming the expression of homozygous Gli3 floxed allele, expression of the cre allele, and the resultant recombined Gli3 (Sup. Fig. 1A). We also confirmed Gli3 knockout in peritoneal macrophages by western blot (Sup. Fig. 1B). Additionally, Gli3 knockout in myeloid cells did not affect overall survival (Sup. Fig. 1C). Using IFA-elicited peritoneal macrophages from *M-Gli3^−/−^* or *WT* mice, we treated cells with 100 ng/ml LPS or DPBS control for 24h and harvested supernatants to quantify inflammatory cytokine levels by ELISA. IRF3 has been reported to regulate CCL2, IL-6 and TNF-α levels, strong regulators of inflammation (38). As expected, LPS stimulation induced a significant increase in the secretion of CCL2, IL-6 and TNFα (Fig. 6C). However, the induction of CCL2 and TNFα secretion was significantly reduced in macrophages from *M-Gli3^−/−^* mice compared with macrophages from *WT* mice (Fig. 6C). This suggests that Gli3 is required for TLR-TRIF-induced cytokine secretion. Interestingly, only basal levels of CCL2 were significantly reduced in the absence of Gli3 suggesting Gli3 regulates both basal and TLR-TRIF-induced levels of CCL2. In contrast, basal TNFα levels were unaffected by Gli3 loss and IL-6 levels were significantly elevated in the absence of Gli3 suggesting Gli3 negatively regulates basal IL-6 levels and positively regulates TLR-TRIF induced IL-6 levels.

## 4. DISCUSSION

The regulation of GLI transcription factors by canonical HH signaling is well established (39). Several studies have shown that the GLI2 family member is regulated in a HH-independent manner (9,10,40). Additional research shows GLI3’s role in the p53 and AKT pathway (41,42) suggesting GLI3’s potential to regulate pathways outside of HH signaling as well. Here, we show that GLI3 is a downstream target of TLR signaling where the TRIF-IRF3 axis directly regulates GLI3 expression (Fig. 6B). TLRs are important immune receptors which are crucial for the innate immune response to pathogens (28,43). TLRs are expressed on a variety of immune and non-immune cells including monocytes. We found a significant induction in GLI3 expression upon challenge of monocytes with LPS (Figure 1). This was found to be due to activation of TRIF and subsequently IRF3 (Figure 3-5). The role of TRIF in regulating GLI3 was validated using the TLR3 ligand Poly(I:C) (which exclusively signals through TRIF) and the TLR4 ligand MPLA (which exclusively activates TRIF but not MyD88) (29). Therefore, we have identified GLI3 as a novel target of TLR-TRIF-IRF3 signaling in monocytes.

Due to its attributes as transcription factor, we hypothesized that TLR-induced Gli3 regulates inflammatory cellular responses. To investigate the role of Gli3 in regulating inflammatory gene expression, we generated *M-Gli3^−/−^* mice, which lack Gli3 expression in myeloid cells such as macrophages. Using IFA-elicited peritoneal macrophages from *M-Gli3^−/−^* and *WT* littermates, we identified a reduction in LPS-induced CCL2 and TNFα secretion in the absence of Gli3. This suggests a major regulatory role of Gli3 in inflammation.

Several studies have suggested a role for GLI3 in the immune system, spanning both innate and adaptive immunity (44,45). GLI3 was found to regulate B- and T-cell development as well as the expression of CD155 on NK cells (44,46,47). More interestingly, in a cDNA microarray analysis, Gli3 was one of the upregulated genes when bone marrow derived macrophages and RAW264 cells were challenged with LPS (48). Therefore, our findings in human monocytes are consistent with studies in mouse macrophages showing an induction of Gli3 in response to LPS stimulation. We characterize the signaling pathway downstream of LPS/TLR4 that regulates GLI3 (Figures 3-5) and identify IRF3 signaling to regulate GLI3. Using macrophages from mice with conditional deletion of Gli3 in myeloid cells (*M-Gli3^−/−^*), we show that Gli3 modulates LPS-induced IL-6, CCL2 and TNF-α levels (Figure 6C). Due to the potency of these inflammatory cytokines, a role of Gli3 in inflammation can be extrapolated. Since Gli3 knockout resulted in a reduction in CCL2 secretion by macrophages, Gli3 may play a role in immune cell trafficking and therefore in the initiation of the inflammatory response. High levels of CCL2 have also been shown to correlate with high cancer occurrence in several cancer types (49–51), suggesting Gli3 may play a role in cancer in addition to pathogen-induced inflammation. Indeed, a role for GLI3 in cancer progression has been suggested (19). Additionally, elevated serum CCL2 levels were associated with multiple sclerosis (MS), rheumatoid arthritis (RA) and atherosclerosis (50,52,53). TNF-α is another inflammatory cytokine that was reported to regulate RA, psoriasis and inflammatory bowl disease (54–56). Since Gli3 knockout reduced LPS-induced TNF-a secretion, Gli3 may modulate the pathology of these diseases. In addition to CCL2 and TNF-α, we show that Gli3 regulates IL-6 secretion (Figure 6C). IL-6 plays a role in the initiation and progression of inflammation. Therefore, it is not surprising that IL-6 plays a role in a multitude of inflammatory diseases. High levels of IL-6 were reported in Castleman’s and Crohn’s disease as well as in RA patients (57,58), suggesting GLI3 may play a role in these diseases. Additionally, increased expression of TLR3 and TLR4 was reported in RA patients and TLR4 was upregulated in Crohn’s disease (59,60), suggesting GLI3 may regulate the pathology of Crohn’s disease and RA through modulation of IL-6.

Upon stimulation of TLR4, signaling can occur through either MYD88 or TRIF adapter proteins (28,61). We identified TRIF as the adaptor protein mediating TLR4 signaling to regulate GLI3 expression in both our cell line models and in primary CD14^+^ cells from PBMCs. Additionally, our data suggests a negative regulation of GLI3 by MYD88, since knockdown of MYD88 using RNAi results in an increase in GLI3 protein expression (Fig. 3A). Downstream of MYD88, NF-κB and MAPK signaling can be activated (26). While targeting MAPK with pharmacological inhibitors did not change GLI3 levels, inhibition of NF-κB resulted in an increase in GLI3 protein expression (Figure 4). This suggests a potential role for MYD88 as a negative regulator of GLI3 by activating NF-κB to decrease GLI3 levels. MYD88 is known to be a protein regulating inflammatory cytokine production while TRIF-IRF3 signaling is known to regulate the production of interferons in addition to proinflammatory cytokines (62). Siednienko et al (2011) suggest a negative regulatory effect of MYD88 on interferon production since the absence of MYD88 increased TLR3-dependent phosphorylation of IRF3 (63). Therefore, it can be extrapolated that MYD88 negatively regulates GLI3 by compromising IRF3 phosphorylation. However, it remains unclear what role NF-κB plays in this inhibitory effect and future studies focused on analyzing the role of NF-κB in MYD88-dependent inhibition of GLI3 would increase our understanding of this potential negative feedback signaling.

Several transcription factors were shown to regulate the promoter of GLI1 and/or GLI2 (9,10,40,64,65). However, our understanding of the regulation of the GLI3 promoter is significantly lacking. The human GLI3 gene is composed of 15 exons with transcription start signal (ATG) located in exon 2. Here we examined approximately 3000 bp sequences upstream of exon 1 and exon 2 and identify candidate IRF3 consensus sequences upstream of exon 2 (which contains ATG signal) (Figure 5). In our cell lines, IRF3 binds to the GLI3 promoter region and this binding is enhanced upon TLR-TRIF stimulation (Fig. 5A & 5B). However, in resting CD14^+^ cells from PBMCs, IRF3 does not bind the GLI3 promoter unless TLR-TRIF signaling is activated (Fig. 5C). Therefore, these studies identify a novel TF (IRF3) that regulates GLI3 by binding to the GLI3 promoter region.

In summary, we identified TLR signaling as novel HH-independent signaling pathway that regulates GLI3 and showed this regulation is dependent on TRIF-IRF3 signaling axis. Additionally, to our knowledge, this is the first report of the regulatory components of the TLR-GLI3 axis where IRF3 was identified as a novel transcription factor that can modulate the promoter region of GLI3. These results increase our understanding of the signaling molecules that can regulate GLI3 in a HH-independent manner. Furthermore, we show that Gli3 is required for TLR-induced regulation of CCL2 and TNFα, therefore identifying a novel role for Gli3 in mediating TLR-induced inflammation.

## ACKNOWLEDGEMENTS

This research was supported by an NIH COBRE Center of Integrated Biomedical and Bioengineering Research (CIBBR, P20 GM113131) through an Institutional Development Award (IDeA) from the National Institute of General Medical Sciences.

## AUTHORSHIP

S.J.M. performed experiments and wrote the paper; W.H., A.H and M.K. performed experiments; R.R. provided reagents and professional conceptual insight; and S.F.E. conceptualized the study, wrote the paper and oversaw all aspects of the research.

