## Supplementary figures and images for "A novel mechanism of regulation of the transcription factor GLI3 by toll-like receptor signaling"

### Supplemental Figure 1

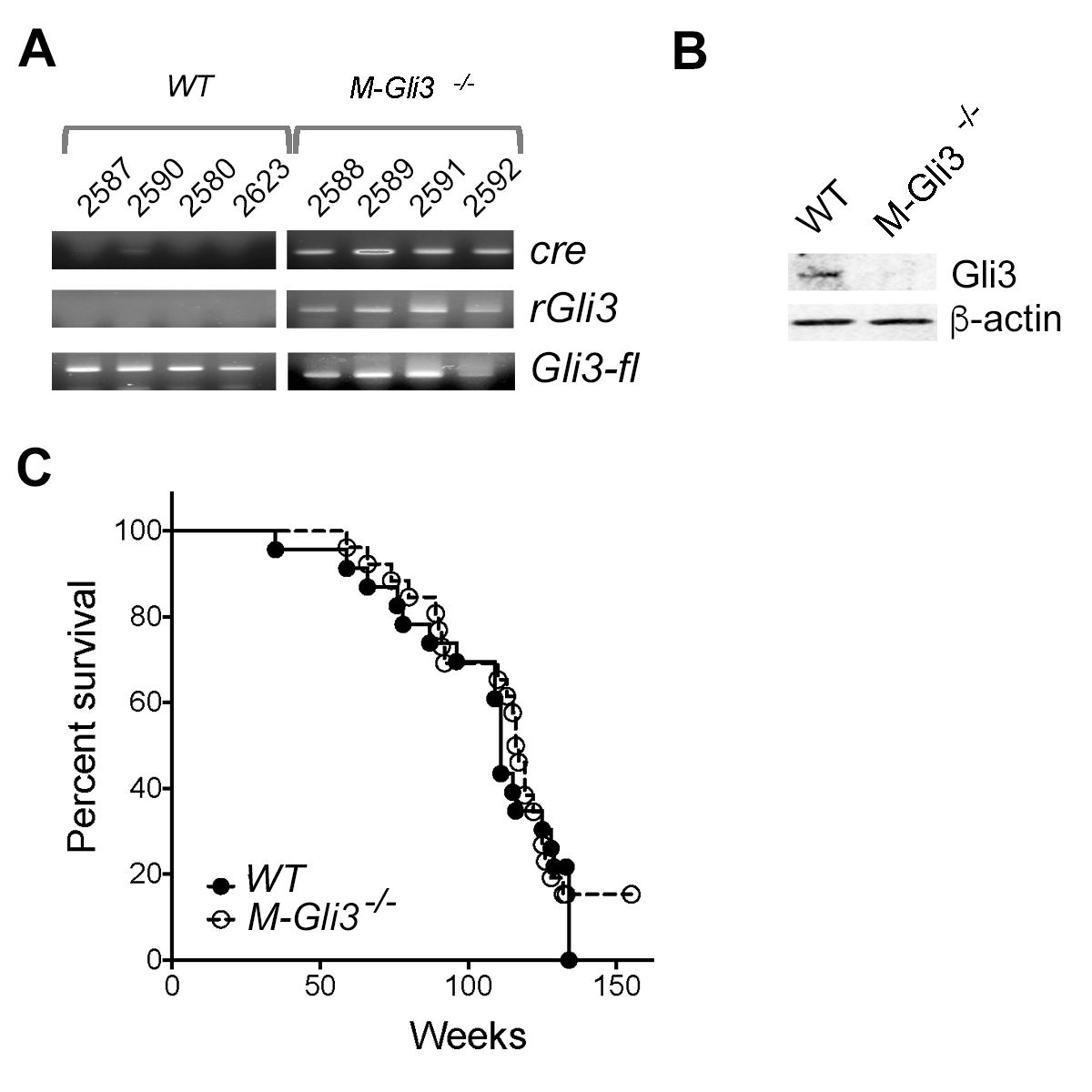
